# Conditional filamentation enhances bacterial survival in toxic environments

**DOI:** 10.1101/2025.05.13.653778

**Authors:** O.B. Aguilar-Luviano, F. Santos-Escobar, S. Orozco-Barrera, R. Peña-Miller

**Affiliations:** Programa de Biología de Sistemas, Centro de Ciencias Genómicas, Universidad Nacional Autónoma de México, 62210, Cuernavaca, México; Departamento de Biología, División de Ciencias Naturales y Exactas, Universidad de Guanajuato, 36050, Guanajuato, México

**Author notes:** These authors contributed equally (O.B. Aguilar-Luviano, F. Santos-Escobar).

**Keywords:** Bacterial filamentation, Phenotypic plasticity, Transient resistance

## Abstract

Bacterial phenotypic plasticity enables rapid adaptation to fluctuating environments. Filamentation, a shape-shifting response commonly observed under stress, has often been viewed as a byproduct of cellular damage. However, filamentation might also confer survival advantages by influencing toxin accumulation dynamics. In this study, we examined the adaptive value of filamentation in *Escherichia coli* using a genetically controlled, SOS-independent induction system to compare isogenic, yet phenotypically distinct cells. By integrating mathematical modeling, single-cell microfluidics, and time-resolved flow cytometry, we evaluate bacterial survival under heavy metal and *β* -lactam antibiotic stress. Our results show that filamentation can improve survival by decreasing the surface area-to-volume ratio, which slows intracellular toxin accumulation and extends the time available for stress response activation or for external toxin levels to dissipate. These findings suggest that filamentation serves as an effective morphological strategy to transiently withstand environmental toxicity, reinforcing its broader role in bacterial stress adaptation.

## 1 Introduction

Bacteria adjust their shape in response to stressful environments through morphological plasticity.^1, 2^ Filamentation—the elongation of cell length—is a response commonly observed under antibiotic and heavy metal stress.^3^ While often interpreted as a byproduct of cellular damage, recent studies have suggested that filamentation may actively influence bacterial resilience by affecting the dynamics of toxin accumulation.^4, 5^ In pathogenic bacteria, filamentation has been shown to enhance survival by protecting against phagocytosis and contributing to biofilm formation, which can increase antimicrobial resistance.^6, 7^ This morphological adaptation, triggered by factors such as DNA damage^5^ and inhibition of cell wall synthesis,^8^ allows bacteria to endure stress, potentially increasing their chances of survival until conditions improve.

Beyond its role in stress tolerance, conditional filamentation supports surface colonization^9^ and aids immune evasion.^9–11^ This morphological adaptation is widespread across bacterial species in both environmental and host-associated contexts,^12–14^ influencing bacterial competition,^15^ persistence within host tissues,^16^ and cell-to-cell spreading between hosts.^17^ Filamentation also contributes to broader ecological interactions, including adaptations in soil communities^18, 19^ and colony-scale behaviors such as chirality^20^ and growth under physical confinement.^21^

Bacterial filamentation is directly regulated by the SOS response, a well-characterized bacterial stress-response mechanism that increases resistance to diverse environmental stressors, including heavy metals^22, 23^ and antimicrobial substances.^24, 25^ This response involves the upregulation of numerous genes, including those involved in DNA repair (e.g., *recA, lexA*), mutagenesis (e.g., *umuC, dinB*), and filamentation (e.g., *sulA*), which collectively enhance bacterial survival under adverse conditions. SulA, for instance, inhibits FtsZ assembly by sequestering monomers, thereby preventing Z-ring formation and halting cell division.^26^ The SOS system is also known to promote genetic variation through mutagenesis,^27, 28^ facilitate the horizontal transmission of virulence factors,^29^ and promote the spread of antibiotic resistance genes via conjugation.^30, 31^ Additionally, the SOS response is necessary to trigger gene cassette shuffling in mobile integrons, a mechanism that enhances bacterial adaptability by rearranging and expressing different genes in response to environmental stress.^32^

Induction of the SOS response can be triggered by various stressors that disrupt DNA replication and cell division processes^33^ (Figure 1A). Stress-response systems that monitor and repair cell wall perturbations, such as those caused by bactericidal antibiotics, play a critical role in bacterial survival and adaptation.^34^ For instance, *β* -lactam antibiotics like ampicillin induce the SOS response by disrupting cell wall synthesis, leading to filamentation and enhanced tolerance to cell wall damage.^1, 14, 35^ Similarly, heavy metals can induce the SOS response through DNA damage and oxidative stress, activating pathways involved in DNA repair.^36^ Although SOS-induced filamentation delays cell division during periods of stress, it represents just one component within the broader and complex SOS regulatory network.^37, 38^ The regulatory complexity of the SOS system complicates efforts to isolate the specific contribution of cell morphology to transient resistance, highlighting the importance of distinguishing shape-driven effects from other SOS-mediated processes.

**Figure 1.**
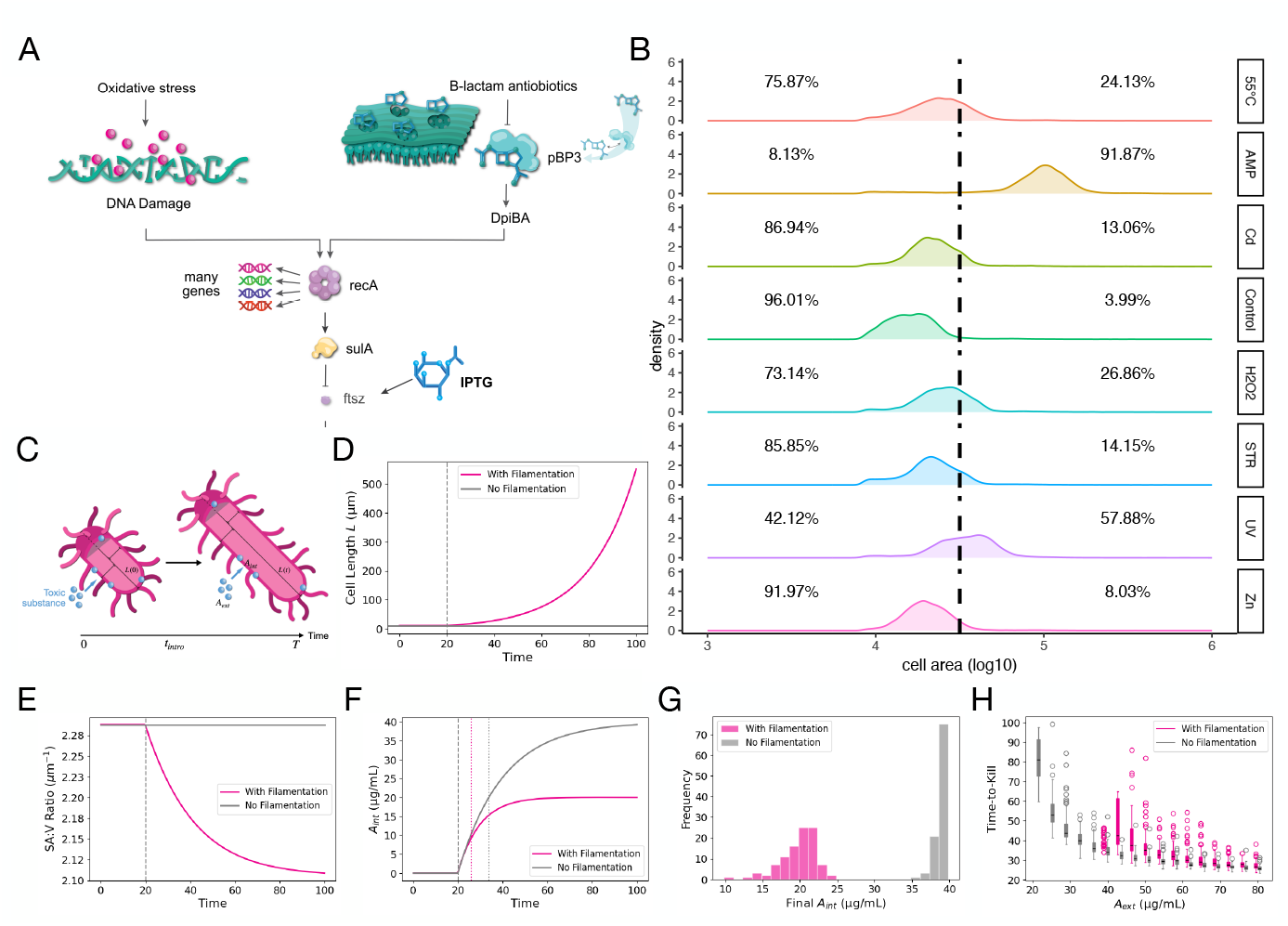
Modeling the impact of filamentation on toxin accumulation and survival. (A) Diagram illustrating filamentation triggered by DNA damage (oxidative stress) or *β* -lactam antibiotics. The latter activate the DpiBA system; DpiA binds replication forks, exposing single-stranded DNA that recruits RecA, initiating the SOS response. *sulA* expression inhibits FtsZ-ring assembly, blocking cell division and promoting filamentation. The synthetic system used in this study employs an inducible promoter with IPTG regulation of *ftsZ* (strain VIP205). (B) Cell size distribution under multiple stressors, each represented by a different color. The dotted line indicates the threshold for classifying cells as filamented, set at the 95th percentile of cell sizes in the control group. (C) Schematic representation of the mathematical model, illustrating antibiotic uptake and accumulation in a filamenting bacterial cell. The cell is modeled as a rod with hemispherical caps, where the length *L*(*t*) is a time-dependent variable. As *L*(*t*) changes, the periplasmic volume (*V*_peri_) and periplasmic surface area (*SA*_peri_) are consequently adjusted. The external antibiotic concentration (*A*_ext_) represents the drug concentration in the surrounding environment, while the internal antibiotic concentration (*A*_int_) depends on *A*_ext_ and varies over time as antibiotics diffuse into the periplasmic space. (D) Cell length over time in filamenting and non-filamenting cases. In the filamentation case, *L* increases following antibiotic exposure, whereas in the control case, *L* remains constant. (E) Surface area-to-volume ratio (SA/V) over time (*µ*m^−1^). Filamentation leads to a progressive decrease in SA/V, reducing antibiotic accumulation per unit volume. (F) Antibiotic concentration (*A*_int_) in the periplasm as a function of time, with lethal thresholds indicated with dotted lines. In filamenting cells, *A*_int_ reaches lethal levels more slowly than in non-filamenting cells, illustrating the protective effect of reduced SA/V. (G) Histogram of final periplasmic antibiotic concentrations (*A*_int_) in filamenting and non-filamenting populations. Variability in antibiotic uptake was introduced by sampling the influx parameter (*k*_in_) from a distribution, leading to a broader range of *A*_int_ values in filamenting cells. (H) Dose-response analysis of time-to-kill with and without filamentation shows that filamentation extends time-to-kill at higher antibiotic concentrations, reflecting the delayed intracellular accumulation of toxins, and prevents lethal accumulation at lower concentrations.

In this study, we aim to clarify the specific adaptive role of filamentation by isolating it from the SOS response using an *Escherichia coli* strain engineered with an inducible promoter controlling the cell division gene *ftsZ*.^39^ By decoupling filamentation from the canonical SOS response, we can directly test whether morphological changes alone improve bacterial survival under toxic stress conditions. Using this controlled experimental system combined with mathematical modeling, single-cell microfluidics, and time-resolved flow cytometry, we evaluate how filamentation influences survival under heavy metal and *β* -lactam antibiotic exposure. Our results indicate that filamentation enhances bacterial survival by decreasing intracellular toxin accumulation through a reduced surface area-to-volume ratio. These findings support filamentation as an adaptive morphological strategy that allows bacteria to transiently withstand environmental stress. More broadly, the ability of diverse bacterial species to filament in response to multiple stressors suggests that morphological plasticity provides a versatile survival advantage across fluctuating environments.

## 2. Results

### 2.1 Filamentation as a General Bacterial Stress Response

Filamentation has been observed across diverse bacterial species and environmental conditions, suggesting it may function as a general stress response. To test this hypothesis in *Escherichia coli*, we first examined whether filamentation is triggered by a broad range of environmental stressors that disrupt different cellular processes. Using flow cytometry and batch cultures, we assessed filamentation under multiple toxic conditions. Mid-exponential phase cultures were exposed to lethal concentrations of various toxic agents for one hour, including UV radiation (254 nm, 30 J/m^2^), cadmium chloride (CdCl_2_, 50 *µ*M), antibiotics (ampicillin, 20 *µ*g/mL; streptomycin, 50 *µ*g/mL), hydrogen peroxide (2 mM) as a source of reactive oxygen species, zinc sulfate (ZnSO_4_, 100 *µ*M), and heat shock at 55°C.

As illustrated in Figure 1B, all tested stressors induced measurable shifts in the cell size distribution, with elongated cells observed in all conditions relative to the untreated control population. The extent of filamentation varied across stressors, indicating that while filamentation is a widespread response, its magnitude is stress-dependent. We quantified the fraction of filamented cells under each condition. Ampicillin (91.87%) and UV radiation (57.88%) induced the strongest filamentation responses, while oxidative stress (26.86%) and temperature stress (24.13%) also led to substantial increases in cell length. In contrast, streptomycin (14.15%), cadmium (13.06%), and zinc (8.03%) triggered filamentation to a lesser degree.

These findings indicate that filamentation is not exclusively linked to antibiotics targeting cell division but is instead a general response to diverse environmental challenges. Stressors such as ampicillin and UV radiation, which directly interfere with DNA replication or peptidoglycan synthesis, elicited the most pronounced filamentation, consistent with previous reports that filamentation can be a consequence of disrupted cell cycle progression.^40^ In contrast, stressors like oxidative stress and heavy metals, which induce broad cellular damage, triggered filamentation at lower frequencies, possibly through indirect regulatory mechanisms. This variability underscores the complexity of bacterial morphological plasticity and suggests that different stressors activate filamentation via overlapping but distinct pathways.^41, 42^

### 2.2 Mathematical Modeling of Toxic Agent Uptake in Filamenting Bacteria

To evaluate whether bacterial filamentation provides a survival advantage under toxic stress, we used a simple mathematical model to describe how morphological changes influence the intracellular dynamics of toxin accumulation. Therefore model represents bacterial cells as rods with hemispherical caps, where filamentation alters cell length (*L*), and therefore the periplasmic volume (*V*_peri_) and surface area (*SA*_peri_), ultimately affecting the rate of toxin accumulation inside the cell (Figure 1C). We assume uniform toxin influx across the cell surface, passive diffusion as the primary uptake mechanism and no metabolic degradation. Additionally, we assume filamentation affects only cell geometry without altering membrane permeability or other physiological processes.

As bacterial cells elongate, the periplasmic volume increases more rapidly than the surface area, reducing the surface area-to-volume (SA/V) ratio (Figure 1E). Because passive diffusion is proportional to SA/V, this reduction slows toxin influx relative to cell volume, delaying intracellular toxin accumulation and extending the time required to reach the lethal concentration. Numerical simulations performed with parameters described in Table S2 illustrate these effects in both filamenting and non-filamenting conditions. In the filamentation scenario, bacterial cell length (*L*) increased significantly following toxin exposure, while in non-filamented controls, it remains constant (Figure 1D). The resulting reduction in SA/V delays intracellular toxin accumulation, thereby increasing the time window before lethal concentrations are reached (Figure 1F).

Asymmetric partitioning of efflux pumps, such as AcrAB-TolC, has been shown to generate phenotypic heterogeneity in drug resistance, leading to systematic differences in growth and survival under antibiotic stress.^43^ To incorporate cellular heterogeneity, we introduced variability in the influx parameter (*k*_in_), simulating natural differences in membrane permeability or toxin uptake rates across individual cells. The resulting distributions of antibiotic concentrations in the periplasm (Figure 1G) showed greater variability among filamented cells compared to non-filamented cells. We further performed a dose-response analysis to assess how different initial cell lengths and antibiotic influx rates affect the time to cell death (Figure 1H). Our theoretical results indicate that filamentation significantly extends survival time, particularly at higher antibiotic doses, highlighting the protective effect of delayed toxin accumulation. At lower antibiotic doses, intracellular concentrations often fail to reach lethal thresholds within the simulated timeframe, further supporting the hypothesis that filamentation serves as an effective buffer against environmental stress.

### 2.3 Filamentation Reduces Surface Area-to-Volume Ratio and Dilutes Antibiotic Uptake

To test the predictions of our mathematical model, we used a microchemostat device, a microfluidic system that enables long-term imaging of single bacterial cells under precisely controlled environmental conditions.^44^ This setup maintains a continuous flow of fresh medium, supporting stable growth while allowing for rapid environmental shifts such as antibiotic exposure and removal. By tracking individual *E. coli* cells in real time, we quantified how filamentation affects SA/V ratio and intracellular toxin accumulation, as well as observed morphological changes before, during, and after antibiotic exposure (Methods).

Cells were exposed to ampicillin (20 *µ*g/mL), a *β* -lactam antibiotic that induces filamentation by binding to penicillin-binding protein 3 (PBP3), an essential component of septal peptidoglycan synthesis. In *E. coli*, PBP3 inhibition activates the DpiBA two-component system, which upregulates *sulA*, an SOS effector that inhibits septation by preventing FtsZ polymerization.^45^ As a result, cell division is transiently suppressed,^46, 47^ leading to filamentous growth (Figure 2A).

**Figure 2.**
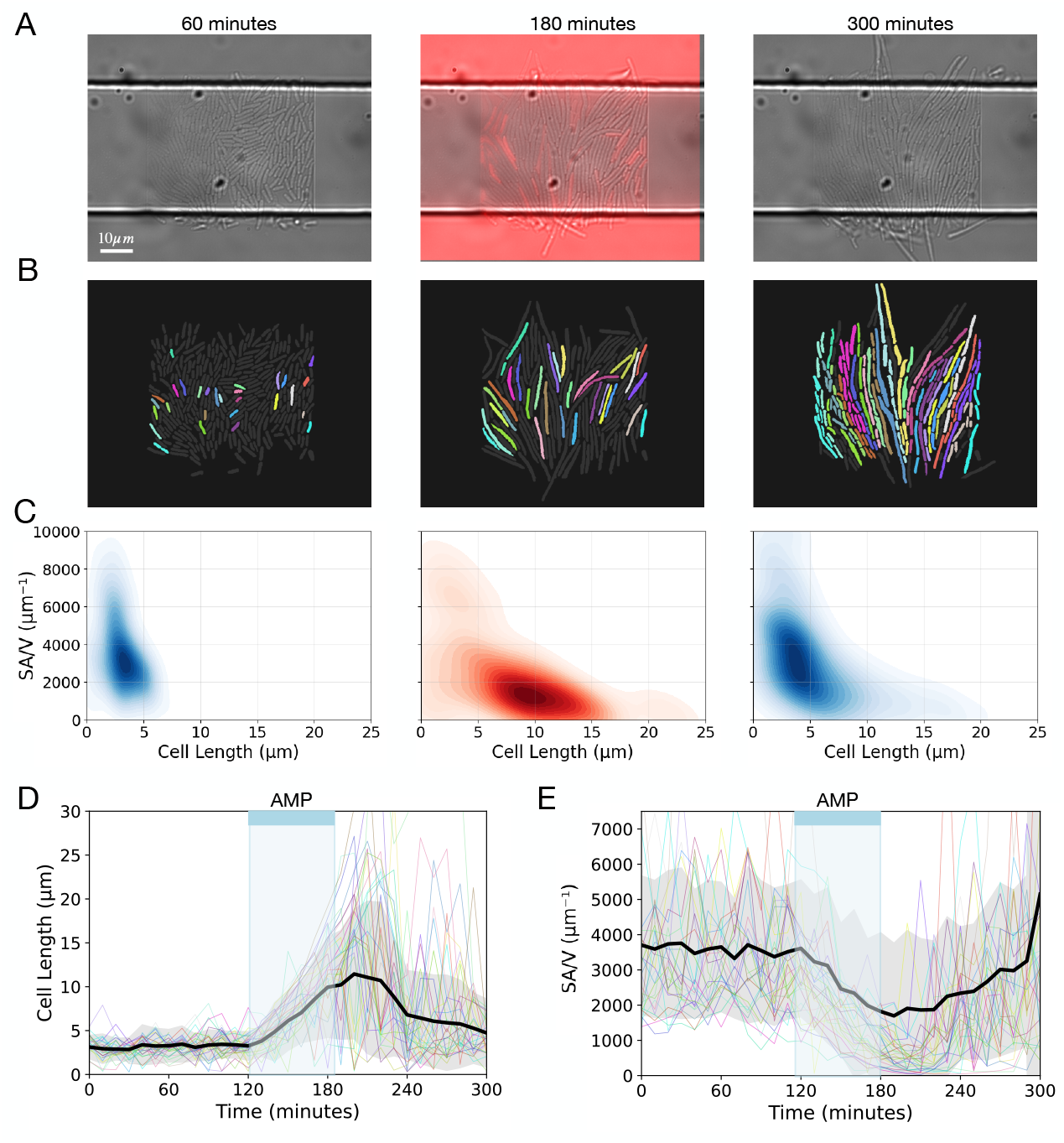
Filamentation reduces surface area-to-volume ratio. **A)** Overlay of phase contrast and red fluorescence images of *E. coli* cells in a microfluidic trap at three time points: before antibiotic exposure (left), during exposure to ampicillin (20 *µ*g/mL) and a red fluorescent dye rhodamine (center), and after antibiotic removal (right). Scale bar: 10 *µ*m. **B)** Cell tracking masks corresponding to (A), with colors representing tracked cells observed throughout the entire experiment. Cells shown in gray are included in the population-level analysis. **C)** Contour plots of cell length (in *µ*m) versus surface area-to-volume (SA/V) ratio (in *µ*m^−1^) at the same time points. During antibiotic exposure (middle, distribution in red), we observe an increase in cell length accompanied by a corresponding decrease in SA/V ratio. **D)** Cell length (µm) over time (minutes). The black line represents the mean population-level cell length, with the gray area indicating the standard deviation. Colored lines show individual cell trajectories corresponding to cells analyzed in panel C. **E)** Surface area-to-volume (SA/V) ratio (µm^−1^) over time (minutes). The black line represents the mean SA/V ratio, with the gray area indicating the standard deviation, with colored lines illustrating single-cell trajectories.

To examine these morphological changes in real time, we tracked individual cells in the microchemostat across the entire antibiotic treatment cycle, capturing dynamics before, during, and after AMP exposure. Initially, cells maintained a consistent rod shape, but upon antibiotic addition, they stopped dividing and began forming filaments. During the AMP pulse, we also introduced rhodamine, a red fluorescent dye used to calibrate our microfluidic system, which allowed us to assess membrane integrity and identify dead cells by red fluorescence. Following AMP removal, surviving cells resumed division and returned to their baseline morphology.

To systematically quantify the effects of AMP on cell morphology, we applied an image analysis pipeline that segmented and tracked bacterial cells from time-lapse movies (Figure 2B). Surface area and volume were estimated using a cylindrical-hemispherical model, allowing us to quantify SA/V dynamics at high temporal resolution (Methods). During antibiotic exposure, we observed a significant increase in mean cell length (3.240 *µ*m to 8.563 *µ*m, *p <* 0.001), accompanied by an increase in both surface area (1.32-fold, p*<* 0.001) and volume (4.26-fold, p *<* 0.001).

Since volume increased faster than surface area, the mean SA/V ratio decreased upon antibiotic exposure (Figure 2C), shifting from 3591.436 *µ*m^−1^ to 1989.202 *µ*m^−1^ (*p <* 0.001). After stress removal, cells resumed division, with mean cell length returning to baseline (Figure 2D) and SA/V reverting to pre-treatment levels (Figure 2E; 3591.436 to 3253.654 *µ*m^−1^, *p >* 0.1). These results confirm that filamentation transiently reduces SA/V during antibiotic exposure, consistent with our model’s prediction that this morphological shift slows intracellular toxin accumulation and delays toxicity onset.

### 2.4 Conditional Induction of Filamentation and Its Impact on Cell Morphology

To decouple filamentation from the SOS response, we used *E. coli* strain VIP205,^48^ which carries an IPTG−inducible promoter controlling the expression of *ftsZ*, a gene essential for septum formation and cell division. In the absence of IPTG, *ftsZ* expression is suppressed, preventing septation and leading to filamentation. When IPTG is added, *ftsZ* expression is restored, allowing normal division (Methods). As a control, we used MC-SCFP3A, a wild-type *E. coli* strain constitutively expressing cyan fluorescent protein (CFP). None of the strains showed statistically significant differences in maximum growth rate under stress-free conditions (p *>* 0.01). VIP205 exhibited a rate of *µ*_*max*_ = 0.864 ± 0.122 h^−1^, compared to *µ*_*max*_ = 1.355 ± 0.022 h^−1^ for the wild-type MC1061 strain, while MC-SCFP3A showed a rate of *µ*_*max*_ = 1.339 ± 0.101 h^−1^.

To examine the effects of filamentation, we used ampicillin (AMP) and cadmium (Cd), which represent two extremes of the filamentation spectrum—AMP induces strong filamentation, while Cd does not (Figure 2B). Dose-response assays confirmed that all strains exhibited comparable stress susceptibilities, with Minimum Inhibitory Concentrations (MICs) of 2 *µ*g/mL for AMP and 0.2 mM for Cd. No statistically significant differences were observed between strains (*p >* 0.1; Figure S1). To induce filamentation, IPTG was removed before AMP or Cd exposure, ensuring that VIP205 cells had already transitioned to the filamentous state before stress application (Figure S2). Cells were then exposed to the toxic agent for 180 minutes, after which the stressor was removed, and IPTG was reintroduced to allow surviving filamented cells to resume division (Figure 3A).

**Figure 3.**
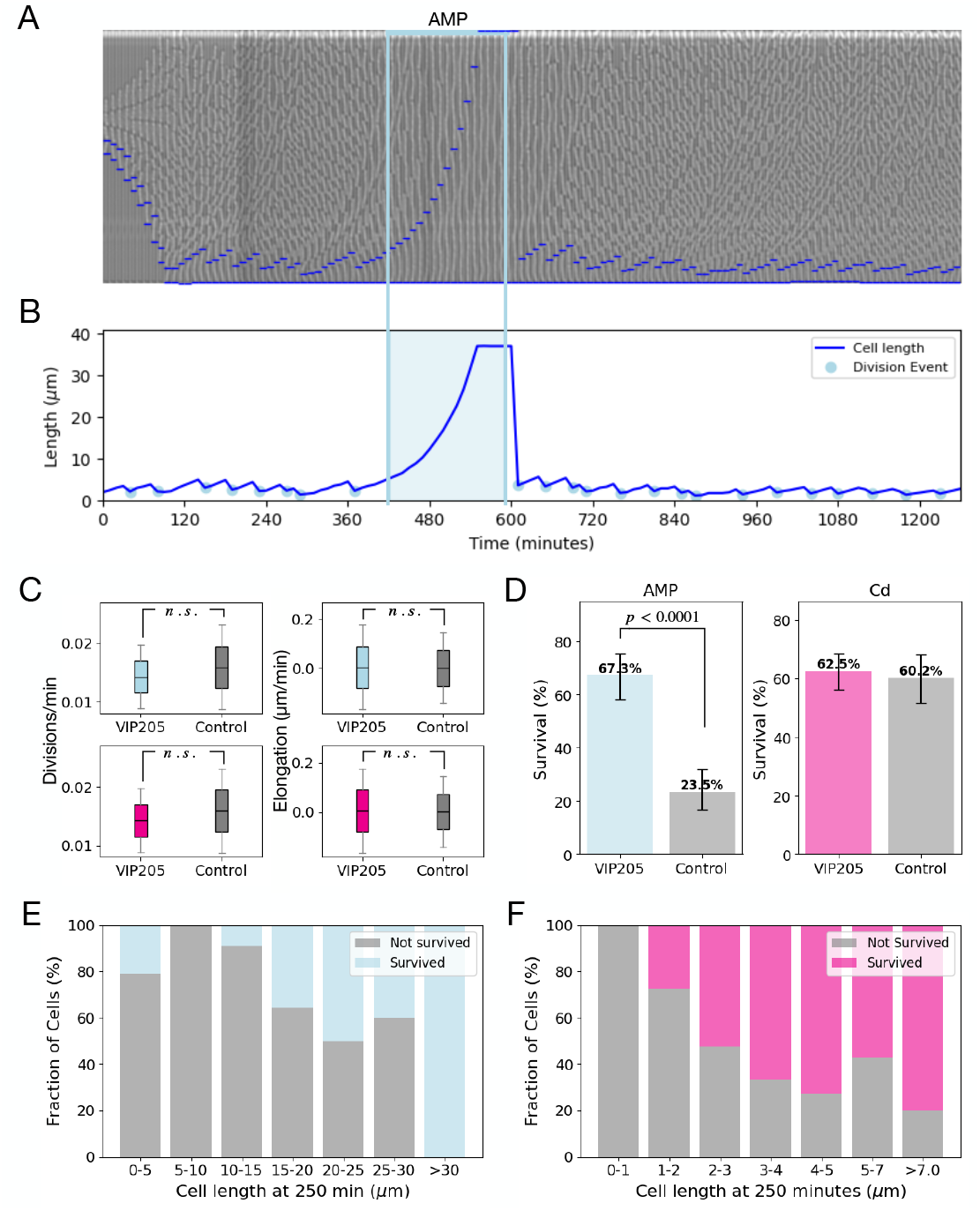
Controlled induction of filamentation and response to toxic stress. **A)** Kymograph showing the time-resolved growth of a mother cell in a microfluidic channel. Time progresses from left to right, with each vertical column representing the same spatial position in the channel at successive time points. The blue horizontal lines mark the boundaries of the tracked cell. The light blue rectangle indicates the period of AMP exposure. **B)** Time-series of cell length (in *µ*m) as a function of time (minutes) for a representative cell. Division events are marked with filled circles and shaded area indicates AMP exposure. **C)** Duplication rates (divisions/min) and average elongation rates (*µ*m/min) of VIP205 and MC-SCFP3A control cells under pre-stress conditions. **D)** Percentage of survival of VIP205 cells exposed to AMP (blue) and Cd (pink). Grey bar illustrates survival of MC-SCFP3A control to the same stress conditions. **E)** Proportion of surviving (blue) and non-surviving (grey) cells grouped by cell length at the time of AMP exposure. **F)** Fraction of surviving (pink) and non-surviving (grey) cells as a function of cell length at the time of Cd exposure.

Furthermore, we tracked single-cell morphology using a dual-input mother-machine microfluidic system^49^ to examine how filamenting and non-filamenting cells responded to toxic stress with hight temporal resolution (Figure 3B; Methods). While the microchemostat captured population-level responses under dynamic conditions, the mother machine enabled precise tracking of individual cell elongation and division over time.^50^ Across three experimental replicates, we examined 232 VIP205 cells and 128 MC-SCFP3A cells exposed to Cd (Figure S3). Similarly, for the AMP experiment, we analyzed a total of 110 VIP205 cells and 119 MC-SCFP3A individual cells for 12 hours (Figure S4).

In AMP-exposed populations, division rates were 0.034 ± 0.0089 divisions/min for VIP205 and 0.039 ± 0.0095 divisions/min for MC-SCFP3A (p*>* 0.1, Mann–Whitney U test, H_0_: equal distributions). Similarly, under Cd exposure, VIP205 and MC-SCFP3A exhibited division rates of 0.0143 ± 0.0055 and 0.0159 ± 0.0072 divisions/min, respectively (p*>* 0.1, Mann–Whitney U test, H_0_: equal distributions). Elongation rates were similar under non-stress conditions (Cd: 0.005 ± 0.0028 *µ*m/min vs. 0.0018 ± 0.0013 *µ*m/min; AMP: 0.034 ± 1.16 *µ*m/min vs. 0.029 ± 0.72 *µ*m/min), with no significant differences between VIP205 and MC-SCFP3A populations (p*>* 0.1, Mann–Whitney U test, H_0_: equal distributions). These results confirm that both strains share comparable baseline growth dynamics, supporting the interpretation that stress-induced differences arise from morphological changes rather than inherent growth variation (Figure 3C).

### 2.5 Filamentation Enhances Survival Independently of the SOS Response

To test whether filamentation provides a survival advantage under toxic stress, we used both single-cell time-lapse microscopy and population-level flow cytometry to compare the fate of filamented and non-filamented populations exposed to AMP and Cd. In the mother machine, filamented cells (VIP205, –IPTG) were more likely to survive and resume growth after stress removal (Figure 3D). Following AMP exposure, VIP205 cells in the induced filamentation condition (IPTG−) showed a significantly higher survival rate (67.3%) compared to MC-SCFP3A cells exposed to the same concentration and duration of stress (IPTG−, 23.5%; *p <* 0.0001). Under Cd exposure, the survival difference between VIP205 and MC-SCFP3A populations was not statistically significant (62.5% vs. 60.2%; *p>* 0.1), likely due to the limited sample size in the mother machine assay.

Importantly, across all conditions tested, cell length was significantly associated with survival. In both AMP- and Cd-exposed populations, surviving cells were consistently longer than non-survivors (Figure 3E-F; Figure S5). For Cd-treated VIP205 populations, the median length of surviving cells after stress removal was 4.84 *µ*m compared to 3.30 *µ*m for non-survivors (Mann–Whitney *U* = 7985.0, *p <* 0.001), while in the CFP control strain, survivors had a median length of 3.41 *µ*m versus 2.31 *µ*m for non-survivors (Mann–Whitney *U* = 2776.0, *p <* 0.001). A similar pattern was observed following AMP exposure. In VIP205, although most cells were filamented due to the SOS response, surviving cells were still longer (median = 36.86 *µ*m) than non-survivors (median = 36.55 *µ*m; Mann–Whitney *U* = 1690.5, *p <* 0.05). In CFP, this difference was more pronounced, with a median length of 29.91 *µ*m in survivors versus 9.73 *µ*m in non-survivors (Mann–Whitney *U* = 1927.0, *p <* 0.001).

To evaluate survival at the population level, batch cultures of filamented (VIP205, IPTG−) and non-filamented (VIP205, IPTG+) cells were exposed to lethal concentrations of AMP or Cd. After exposure, cells were stained with propidium iodide (PI) to distinguish live from dead cells and analyzed using flow cytometry (Figure 4A-B). SSC-Width was used as a proxy for cell length under our experimental conditions, where filamentation leads to increased side scatter. In the non-filamented condition, 83.7% of cells were PI-positive after AMP exposure, whereas in the filamented condition, only 60.5% were PI-positive, indicating increased survival with induced filamentation (Figure 4C). A similar trend was observed under Cd exposure (Figure 4D): non-filamented populations showed 62.8% PI-positive cells, while filamented populations showed a significantly lower fraction (35.0%). In contrast, no-stress controls maintained low PI-positive fractions throughout the experiment (all below 4%), confirming that the observed cell death was stress-induced.

**Figure 4.**
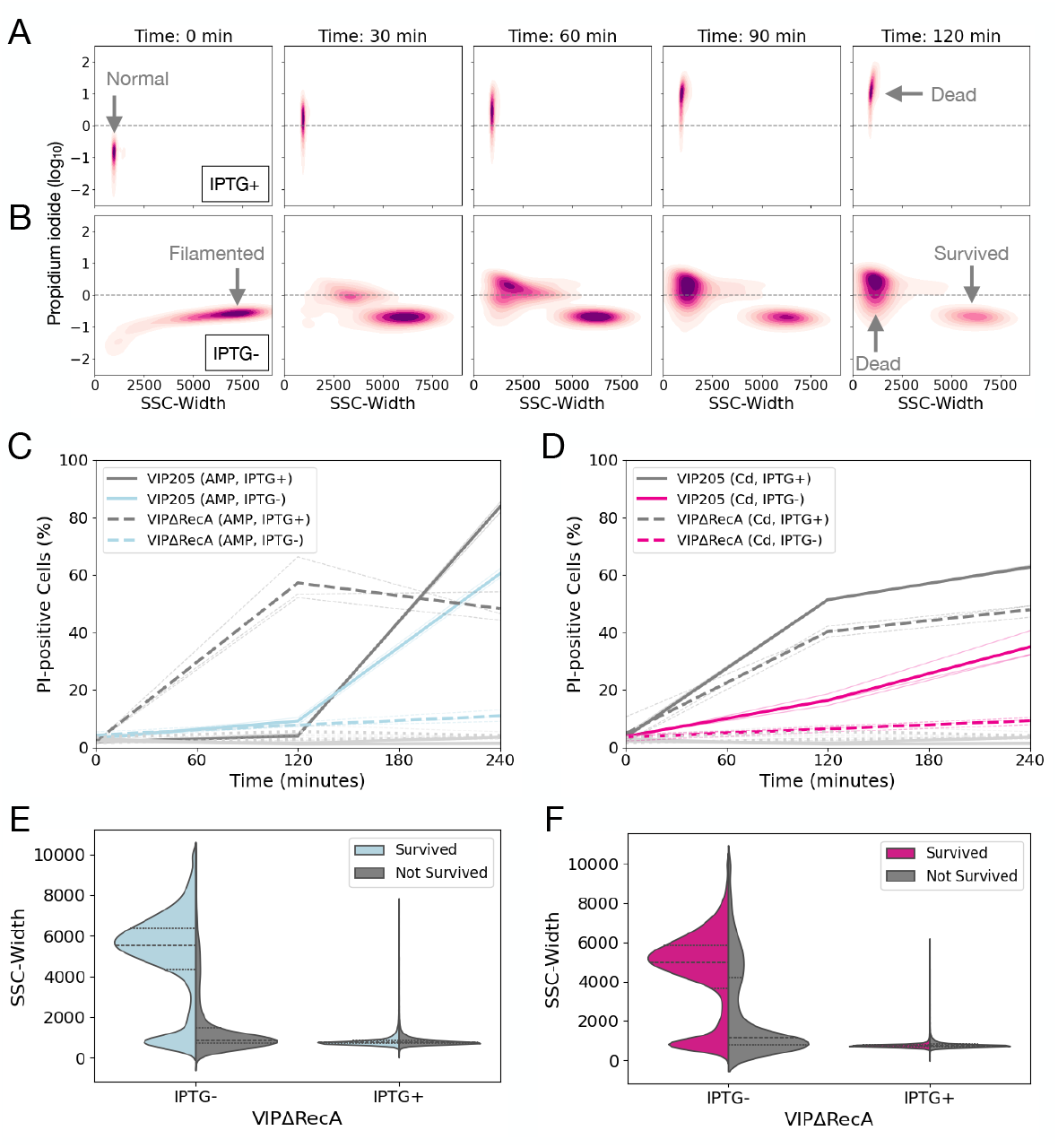
Induced Filamentation Enhances Survival Under Toxic Stress. **A)** Length profiles of sfisl:amentedVcIePldlse(ltVaRIPe2c0A5-+NCods, tIrPeTssG: −) and non-filamented cells (VIP205 + Cd, IPTG+). **B)** Evaluation of cell viability using propidium iodide (PI) staining. **C)** Time-resolved quantification of cell survival under AMP stress. The percentage of PI-positive (non-viable) cells was measured over time for filamented (IPTG−; light blue) and non-filamented (IPTG+; grey) populations. Thick lines represent the mean across replicates; thin lines represent individual experiments. Solid lines indicate strain VIP205; dashed lines indicate strain VIP205 Δ*RecA*. Light grey indicates no-stress controls. **D)** Time-resolved quantification of cell survival under Cd stress. The percentage of PI-positive cells was measured over time for filamented (IPTG−; pink) and non-filamented (IPTG+; grey) populations. Line styles and control coloring are as in (C). **E)** Comparison of cell length distributions between surviving and non-surviving cells after AMP exposure. Violin plots show distributions for filamented (VIP205 Δ*RecA*, IPTG−) and non-filamented (VIP205 Δ*RecA*, IPTG+) cells. Surviving cells are shown in blue, non-surviving cells in grey. **F)** Comparison of cell length distributions between surviving and non-surviving cells after Cd exposure. Conditions and coloring as in (E), with surviving cells shown in pink.

Moreover, consistent with our single-cell data, under both AMP and Cd stress, surviving cells were significantly longer than those that died (p *<* 0.0001 for both conditions). AMP-treated populations also showed signs of filamentation in the IPTG+ condition, suggesting activation of the endogenous SOS pathway (Figure S6). To test whether this effect was RecA-dependent, we repeated the experiment using a VIP205 Δ*RecA* strain^47^ (Table S1). As expected, AMP no longer induced filamentation in the IPTG+ control, whereas the IPTG− condition still triggered filamentation. Surviving cells also showed significantly higher SSC-Widths than non-survivors (AMP: 5015.2 ± 2212.7 vs. 1592.1 ± 1595.4; Cd: 4474.6 ± 2084.9 vs. 2493.2 ± 2298.3; *p <* 0.0001 for both; Figure 4E–F). Altogether, these results show that filamentation promotes bacterial survival under toxic stress, independently of the SOS response.

## 3. Discussion

Bacterial filamentation is a well-documented response to environmental stress, including exposure to antibiotics and heavy metals. This process involves changes in cell morphology that can enhance bacterial survival under adverse conditions.^4^ Filamentation is a conserved response observed across diverse bacterial species and stressors, including nutrient limitation in *Caulobacter crescentus*,^51^ heavy metal exposure in *E. coli, Shewanella oneidensis*, and *Rhodobacter capsulatus*,^12, 13, 22^ and heat or saline stress in *Pseudomonas putida*.^9^ Significant elongation has also been observed in *E. coli* under *β* -lactam antibiotic stress.^14, 52–54^ This widespread occurrence across phylogenetically distinct bacteria and diverse stress types underscores its ubiquity and suggests that filamentation is not an inevitable consequence of cellular damage but a general adaptive strategy to mitigate toxic stress.^6^

In this study, we decoupled filamentation from the SOS response using an inducible expression system controlling *ftsZ*,^48^ and further validated this independence by showing that filamentation-induced survival persists even in a Δ*RecA* background. Single-cell measurements revealed that filamentation delays intracellular toxin accumulation, consistent with predictions from our theoretical model. According to the model, a lower surface area to volume ratio in elongated cells slows down toxin uptake, allowing more time for recovery or adaptation before reaching lethal concentrations. Upon stress removal, filamented cells resumed division and reverted to their original rod-like morphology. These results support filamentation as an active survival strategy that does not rely on SOS-mediated mechanisms.

Additionally, filamentation likely integrates into broader bacterial stress responses, including efflux pumps, porin modifications, and biofilm formation.^55, 56^ By maintaining constant chromosome^1^ and plasmid copy numbers^53^ as cell volume increases, filamented cells ensure that drug-degrading enzymes and efflux pumps are not diluted, allowing them to sustain their activity while intracellular toxin concentrations decrease. Other morphological adaptations have also been implicated in bacterial stress responses. Under cadmium stress, *E. coli* produces minicells that accumulate and expel cadmium sulfide nanoparticles, reducing intracellular toxin levels.^57^ This reinforces the idea that bacterial morphology is not only a passive consequence of stress but an active component of survival strategies.

Recent studies have found metabolic adaptations associated with filamentation that may contribute to bacterial persistence under stress. For instance, filamenting *E. coli* exhibit increased c-di-GMP levels, a secondary messenger involved in biofilm formation and drug tolerance. Inhibiting c-di-GMP synthesis sensitizes cells to *β* -lactams, suggesting metabolic pathways activated during filamentation promote resistance.^58^ These findings further support the idea that filamentation is not an isolated stress response but part of a coordinated survival strategy involving morphological and metabolic adaptations. Further investigation into how filamentation coordinates with the general stress response RpoS, efflux regulation, and other shape-shifting adaptations could clarify its role in bacterial adaptation.^3^

From an applied perspective, investigating the temporal regulation and reversibility of filamentation could identify critical vulnerabilities during morphological transitions, providing potential targets for antimicrobial interventions aimed at disrupting filamentation dynamics. Recent studies have shown that CRISPR-based technologies or chemical inhibitors can be used to destabilize filamentous states, potentially rendering bacteria more susceptible to treatment during periods of acute stress.^59^ Additionally, identifying inhibitors that specifically block morphological transitions—such as septal ring formation or elongation—could help sensitize bacteria to antibiotics. Combining standard antimicrobials with anti-filamentation agents might also enhance therapeutic effectiveness, especially in persistent infections where bacteria rely on morphological adaptations to tolerate treatment. Finally, targeting filamentation could minimize collateral damage to beneficial microbiota, providing a more selective strategy compared to broad-spectrum approaches that rely solely on high antibiotic doses.

Altogether, our results suggest that bacterial filamentation enhances survival under toxic stress by decreasing the surface area-to-volume ratio, thereby slowing intracellular toxin accumulation. Further research into how filamentation integrates with other stress responses, including DNA repair and metabolic adjustments, could reveal additional mechanisms promoting bacterial resilience. From a practical perspective, identifying strategies to selectively disrupt filamentation might enhance the efficacy of antimicrobial therapies, particularly in persistent infections where morphological adaptations confer transient resistance.

## Acknowledgements

We thank A. Fuentes-Hernandez, D. Zamorano, G. Gosset, D. Perez, J.C.R. Hernandez, A. Aertsen and G. Perron for useful discussions. We appreciate A. Fernandez Duque’s contribution in creating the visual illustrations. We also thank Pilar C.P. for assistance with experiments and image analysis. FSE received funding from DGAPA-UNAM. RPM was supported by PAPIIT-UNAM (grant IN209419). OBAL is a student in Programa de Doctorado en Ciencias Bioquímicas, Universidad Nacional Autónoma de México and received fellowship 886346 from CONACYT.

## 4 Materials and Methods

### Strains and Culture Conditions

All *E. coli* strains used in this study are listed in Table S1. The routine growth medium was Luria-Bertani (LB) medium, containing 10 g L^−1^ tryptone, 5 g L^−1^ yeast extract, and 10 g L^−1^ NaCl. When required, antibiotics such as Kanamycin (Kan, 50 *µ*g mL^−1^), Ampicillin (AMP, 20 *µ*g mL^−1^), or Isopropyl *β* -D-1-thiogalactopyranoside (IPTG, 15 or 30 *µ*M) were added to the medium. Liquid cultures were incubated at 37^°^C with vigorous aeration (shaking at 230 rpm).

For cadmium susceptibility assays, a stock solution of cadmium chloride (CdCl_2_, 99.97% purity; SIGMA, USA) was prepared at 100 mM as Cd^2+^ ion in Milli-Q water, filter-sterilized using 0.22 *µ*m syringe filters, and stored at ambient temperature. Working concentrations were prepared by diluting the stock solution with sterile Milli-Q water. To avoid cadmium precipitation during susceptibility analysis, the Heavy Metal MOPS Medium (HMM) was used with slight modifications. HMM was composed of 40 mM MOPS, 50 mM KCl, 10 mM NH_3_Cl, 0.5 mM MgSO_4_, 0.4% glucose, 1 mM glycerol-2-phosphate, 1 mM FeCl_3_, and supplemented with 0.5% Casamino acids (SIGMA). The medium was prepared in Milli-Q water, filter-sterilized using 0.22 *µ*m syringe filters, and stored at 4^°^C. When required, Kanamycin (Kan, 50 *µ*g mL^−1^) or IPTG (10 *µ*M) was added to the HMM. AMP stock solutions (100 mg/ml) were prepared by diluting ampicillin (Sigma-A0166) directly in HMM media.

### Mathematical model

We developed a mathematical model to describe how bacterial filamentation influences the uptake and internal accumulation of a toxic agent. The model assumes that the external concentration, *A*_ext_, remains constant, while the internal concentration, *A*_int_, is uniform within the periplasmic space. The toxic agent enters at a rate proportional to the cell’s surface area. The bacterial cell is modeled as a cylindrical rod with hemispherical caps, where the periplasmic space is defined by an outer radius, *r*_outer_, and an inner radius, *r*_inner_. The periplasmic volume, *V*_peri_(*L*), and surface area, *SA*_peri_(*L*), are functions of cell length, *L*, given by:

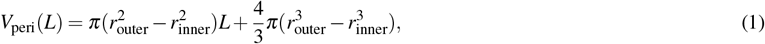

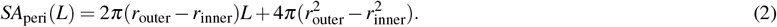

As the cell elongates, the volume and surface area increase at different rates, reducing the surface area-to-volume ratio. This change affects the rate of toxic agent accumulation, effectively diluting the internal concentration over time. The rate of change of *A*_int_ is given by:

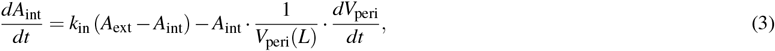

where the first term represents the influx of the toxic agent into the periplasm, and the second term accounts for dilution due to volume expansion. The change in periplasmic volume over time is described by:

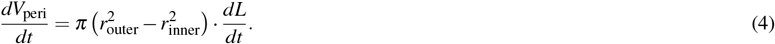

For toxic agents that accumulate in the periplasm, such as *β* -lactam antibiotics, the standard model formulation applies. However, for cytoplasmic toxins like Cd, we set *r*_inner_ = 0, simplifying the volume equation to represent a single intracellular compartment. This modification amplifies the dilution effect, as filamentation increases the total intracellular volume rather than just the periplasmic compartment. In our numerical experiments, filamentation begins at a predefined time point, *t*_intro_, following exposure to the toxic agent. After this point, cell length, *L*, follows an exponential growth model:

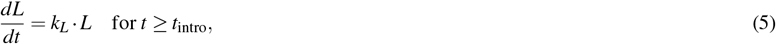

where *k*_*L*_ is the elongation rate constant. In the control (null) model, filamentation does not occur, and the cell length remains constant, allowing comparison between filamenting and non-filamenting cells. Therefore the simulations consist of two phases: pre-exposure (*t < t*_intro_), where no toxic agent enters the cell (*k*_in_ = 0), and exposure (*t* ≥ *t*_intro_), where the toxic agent is introduced and filamentation may occur. Parameter values used in the simulations are listed in Table S2. Simulations were implemented in Python using standard scientific libraries (NumPy, Matplotlib). The ODE system was solved numerically with an adaptive time-step solver. The description of the model and simulation code is available in the project repository (GitHub).

### Characterization of *E. coli* Strain VIP205 in HMM Medium

To maintain normal morphological parameters at high growth rates, *E. coli* VIP205 cultures require the presence of 30 *µ*M IPTG in LB medium.^48^ Since all experiments in this study were conducted in HMM medium, we first determined whether the same IPTG concentration (30 *µ*M) is effective in HMM. An overnight culture of *E. coli* VIP205 grown in LB medium with 30 *µ*M IPTG was washed three times with sterile deionized water, then centrifuged at 3000 x g for 10 minutes and resuspended in HMM medium without IPTG. The culture was then diluted 1:100 in fresh HMM medium without IPTG and incubated at 37^°^C with shaking at 250 rpm for 3 hours. At that point, an aliquot was transferred to a 1% agarose pad to observe filamentous cells, and images were captured using a Nikon Eclipse Ti-E inverted fluorescence microscope with a 100X Plan APO objective (1.45 OIL, Nikon Instruments). Following verification of increased cell length, we restored the normal growth phenotype by adding several concentrations of IPTG (5, 10, 15, and 30 *µ*M). After 3 hours of incubation at 37^°^C and 250 rpm, aliquots were analyzed by microscopy to observe the return of normal growth, as previously described.^48^ Simultaneously, aliquots from each growth stage (normal growth, filamentation induced by IPTG deprivation, and recovery with IPTG addition) were analyzed by flow cytometry using the CYTOFLEX S (Beckman Coulter, USA). For each experiment, 100,000 events were plotted on a log scale, with a threshold on forward scatter (FSC) and side scatter (SSC) set at 200.

### Determination of Cd^2+^ and AMP Toxicity

To evaluate the sensitivity of *E. coli* cells to cadmium (Cd^2+^) and ampicillin (AMP), overnight cultures were diluted to an initial optical density (OD_600_) of approximately 0.05 in fresh LB medium supplemented with kanamycin (Kan, 50 *µ*g/mL) and IPTG (10 *µ*M). Cells were grown in 96-well flat-bottom plates (200 *µ*L per well) at 30°C with continuous shaking in a BioTek Elx808 plate reader (BioTek Instruments, Potton, UK). After 90 minutes of initial growth, cadmium chloride (CdCl2) or ampicillin (AMP) was added to final concentrations ranging from 0 to 200 *µ*g/mL for AMP and from 0 to 0.1 mM for Cd. Experiments were performed in triplicate, with optical densities (OD600) measured every 10 minutes for 8 hours, allowing us to determine maximum growth rates (*µ*_*max*_) and MICs for each stressor.

### Single-cell Microfluidics

We used two microfluidic platforms to continuously observe individual *E. coli* cells under precisely controlled conditions. The first was a microchemostat device^44^ composed confinement chambers that trap approximately 300 − 1, 000 cells in the same focal plane, allowing observation of many cells simultaneously while maintaining constant nutrient supply and facilitating rapid environmental changes. The second a dual input mother machine,^49^ consisting of narrow channels designed to trap single cells, enabling real-time monitoring of growth, morphology, and division behaviors of individual cells over extended periods. Both devices were fabricated were fabricated from polydimethylsiloxane (PDMS) using soft photolithography with SU-8 2000.5 molds (Micro Resist Technology GmbH). The dual-input design consists of comb-like channels that trap mother cells at the bottom, pushing daughter cells into larger channels. Chips were made using Sylgard® 184 Silicone Elastomer at a 1:10 curing agent to base ratio, degassed, and cured at 80°C for 2 hours. After curing, the chips were cut, punched, and bonded to glass coverslips using a plasma cleaner (Harrick PlasmaPDC-001), then baked overnight at 45°C to ensure effective bonding.

For microfluidic experiments, overnight cultures of *E. coli* (VIP205 and MC-SCFP3A strains) were grown in HMM medium with or without 10 *µ*M IPTG, diluted 100-fold into fresh medium, and incubated at 37°C with shaking (230 rpm) until reaching an OD_600_ of ∼0.2-0.3. The cultures were mixed 1:2 (MC-SCFP3A:VIP205), centrifuged, washed with sterile MQ water, and resuspended in fresh HMM medium without IPTG. The dense culture was used to inoculate the microfluidic device. The flow rate was controlled according to Hernandez-Beltran et al. (2020), and cells were allowed to establish baseline conditions before toxic exposure. Growth media was loaded into syringes connected to the PDMS chip via Tygon tubes and Luer connectors. Pressure was controlled with vertical linear actuators and a digital signal generator to maintain precise extracellular conditions. This setup allowed for rapid introduction of toxic agents and environmental switches to observe dynamic bacterial responses.

### Time-lapse image acquisition

High-resolution imaging was performed using a Nikon Eclipse Ti-E epifluorescent microscope equipped with differential interference contrast (DIC), a motorized stage, and a perfect focus system for long-term time-lapse imaging. The microscope was controlled by Nikon NIS-Elements AR software and maintained at 30^°^C using a Lexan Enclosure Unit with Oko-touch temperature control. Time-lapse movies were captured with a 100X Plan APO objective, with no analog gain applied, and field and aperture diaphragms minimized to prevent photobleaching. DIC images were acquired at 9V DIA-lamp intensity with a 200 ms exposure time. For fluorescence imaging, red channel images (excitation 560-590 nm, emission 610-650 nm) were acquired with a 600 ms exposure, and cyan channel images (excitation 430-470 nm, emission 480-520 nm) were acquired with a 200 ms exposure. Images were acquired at 5-minute intervals to ensure high temporal resolution. The microscope was maintained at 30^°^C using a Lexan Enclosure Unit with Oko-touch temperature control.

### Single-Cell Segmentation and Tracking

Segmentation and tracking of bacterial cells in time-lapse movies were performed using a combination of machine learning-based classification, deep-learning-assisted boundary refinement, and manual correction to improve accuracy. Trainable Weka Segmentation (TWS),^60^ a Fiji plugin that applies machine learning algorithms for pixel-based classification, was used to generate probability maps for bacterial cell boundaries based on user-defined training datasets. These probability maps were then processed with CellPose3,^61^ a deep-learning-based segmentation tool, to refine cell contours and improve boundary detection. Following segmentation, cells were tracked using a nearest-neighbor linking approach, enabling the reconstruction of single-cell lineages over time. Morphological parameters, including cell length, width, and fluorescence intensity, were quantified at each frame. Cell volume was estimated using a geometric model that approximates bacterial morphology as a cylinder with hemispherical caps. To determine cell dimensions, each segmented mask was skeletonized to extract its medial axis, ensuring that the cell length measurement followed its true longitudinal shape. The skeleton was then smoothed using a Gaussian filter and a Savitzky-Golay filter, and a B-spline curve was fitted to further refine the centerline. The major axis length was determined from the final smoothed skeleton, while the minor axis was measured perpendicular to the centerline at its midpoint. Using these dimensions, volume was computed as the sum of a cylindrical body and two hemispherical caps.

Image processing of mother machine experiments was performed using ImageJ, from acquisition to producing the kymographs. ImageJ was used to crop and generate kymographs from the time-lapse movies captured during the experiments.Segmentation and tracking of individual cells were performed using a Python-based tool. This bespoke tool was designed for precise tracking and analysis of cells within kymographs. It allows for the interactive setting and correction of the spaces between cells in microfluidic channels, which we refer to as breakpoints. This enables the determination of cell length in consecutive frames and supports analysis of montages from two different channels (e.g., phase and fluorescent). The tracking algorithm iteratively calculates the optimal y-displacement between consecutive frames by comparing small windows of pixels. This algorithm tracks the vertical position of features across frames, adjusting for displacements to maintain accurate tracking. The tool emphasizes interactivity and manual correction to ensure high precision in the analysis. The image analysis code is available in the public [repository](https://github.com/ccg-esb-lab/vip205).

### Time-resolved flow cytometry

Population-level survival assays were performed to evaluate the effect of filamentation on bacterial resilience under toxic stress. Cultures of *E. coli*, induced or suppressed for filamentation via IPTG manipulation, were exposed to a range of toxic agents. Following exposure, cultures were washed with PBS to remove the agents and resuspended in fresh LB medium for recovery. Survival was quantified by plating on LB agar without IPTG, with colony-forming units (CFUs) counted after 24 hours of incubation. Flow cytometry was used to assess cell size distribution and quantify the survival advantage of filamented cells under toxic stress. The survival of filamented and non-filamented *E. coli* populations was compared under lethal doses of cadmium and other agents. Forward and side scatter profiles were analyzed to determine cell viability and distinguish between filamented and non-filamented cells. Flow cytometry was performed using a CytoFLEX S (Beckman Coulter) with detectors for forward scatter (FSC) and side scatter (SSC). Instrument settings included thresholds of 200 for both FSC and SSC, with laser excitation at 488 nm. Data were visualized as dot plots (SSC-H vs. SSC-W) to assess cell size distribution and to distinguish between filamented and non-filamented populations.

## Supplementary material

**Figure S1.**
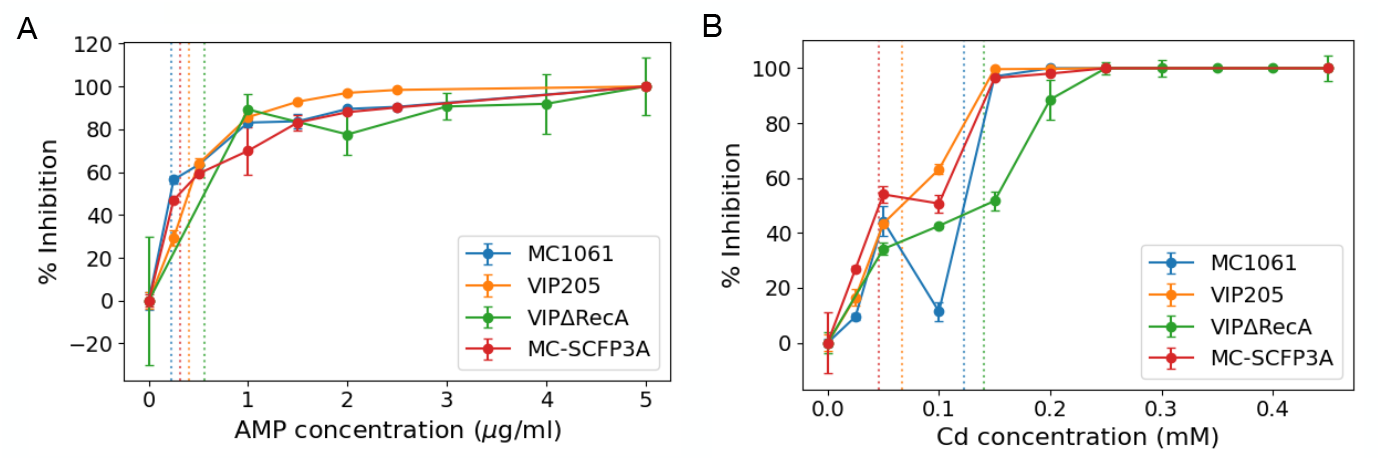
Susceptibility curves showing percentage of growth inhibition at 20 hours as a function of stressor concentration. Lines correspond to strain VIP205 (orange), MC-SCFP3A (green), and the wild-type control MC1061 (blue). **A)** Dose-response to ampicillin (AMP) at increasing concentrations (*µ*g/mL). **B)** Dose-response to cadmium (Cd) at increasing concentrations (mM).

**Figure S2.**
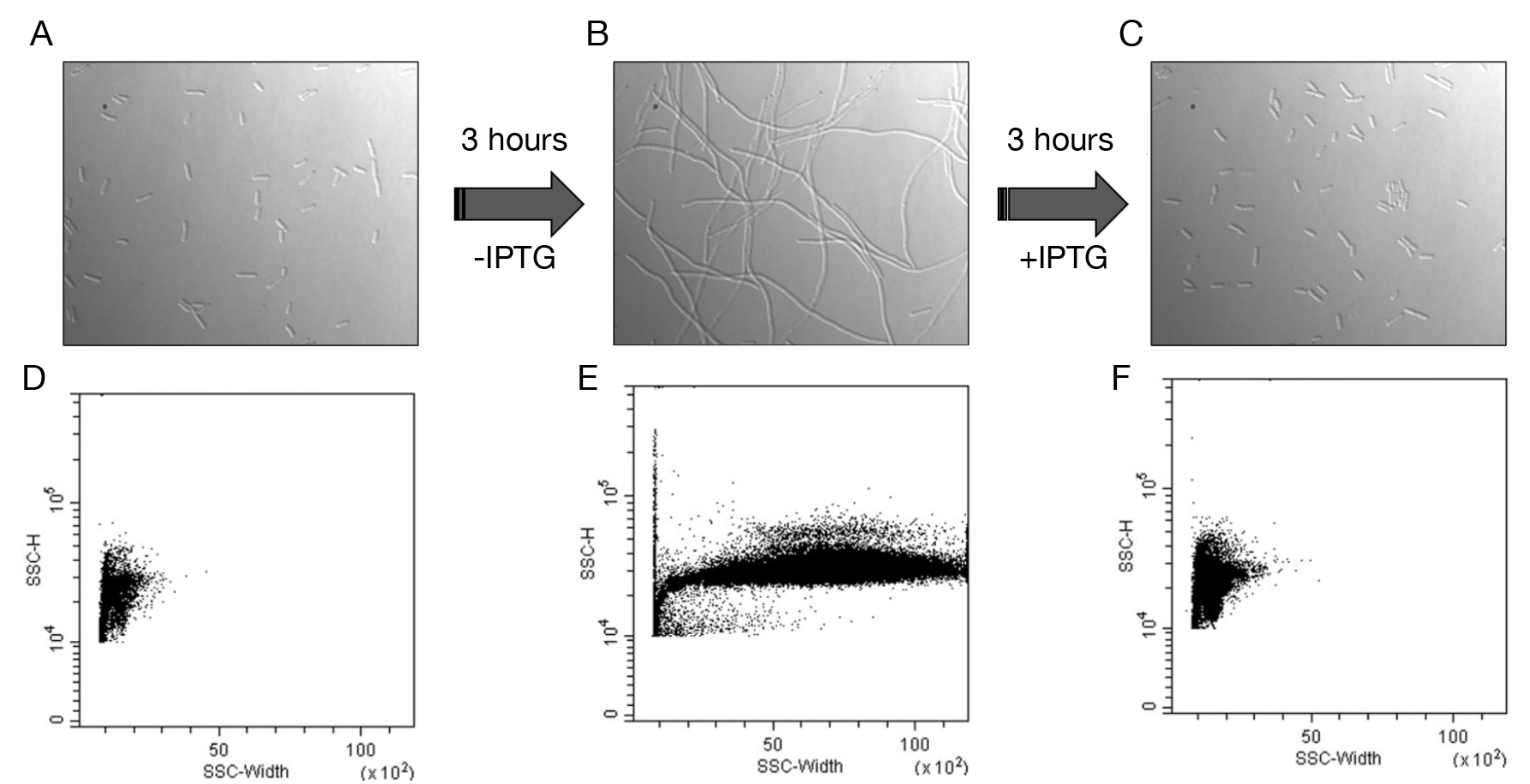
**A)** Microscopy image of *E. coli* VIP205 under normal growth conditions with IPTG (10 *µ*M), showing typical morphology. **B)** Without IPTG, cells exhibit significant elongation, demonstrating filamentation. **C)** Reintroducing IPTG restores normal morphology. **(D-F)** Flow cytometry scatter plots for VIP205 cells. **D)** Normal cell size distribution with IPTG. **E)** Absence of IPTG shifts the plot rightward, reflecting elongation. **F)** IPTG reintroduction reverts the distribution to baseline sizes.

**Figure S3.**
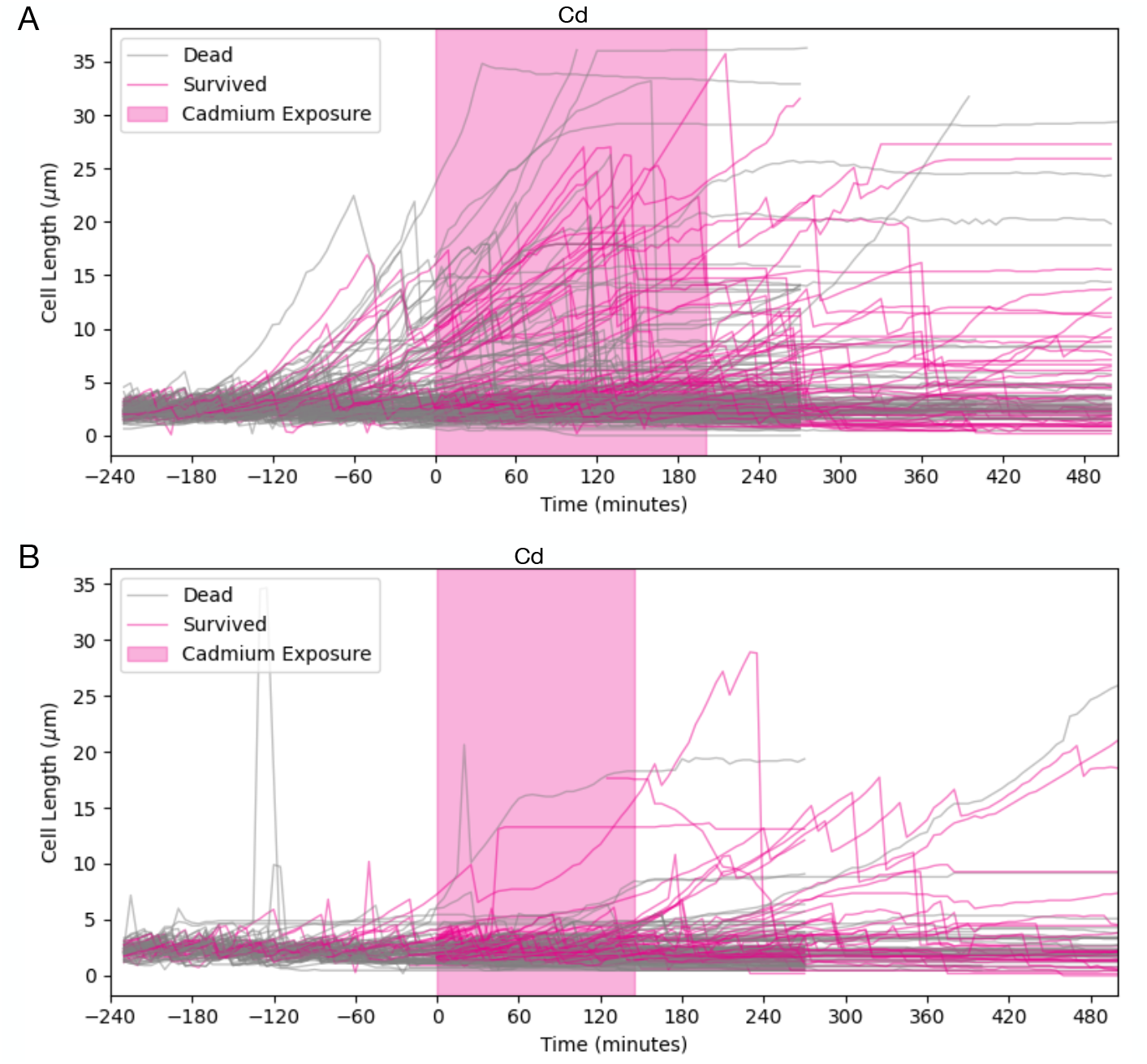
**A)** Time-series of individual cell lengths (*µm*) for VIP205 cells in the mother machine. The pink shaded region indicates the period of cadmium (Cd) exposure. Surviving cells, which resumed division after Cd removal, are shown in pink, while non-survivors appear in gray. **B)** Time-series of individual cell lengths (*µm*) for control (MC-SCFP3A) cells in the mother machine. The Cd exposure period is shaded in pink. Cells that survived are shown in pink, while those that did not are in gray.

**Figure S4.**
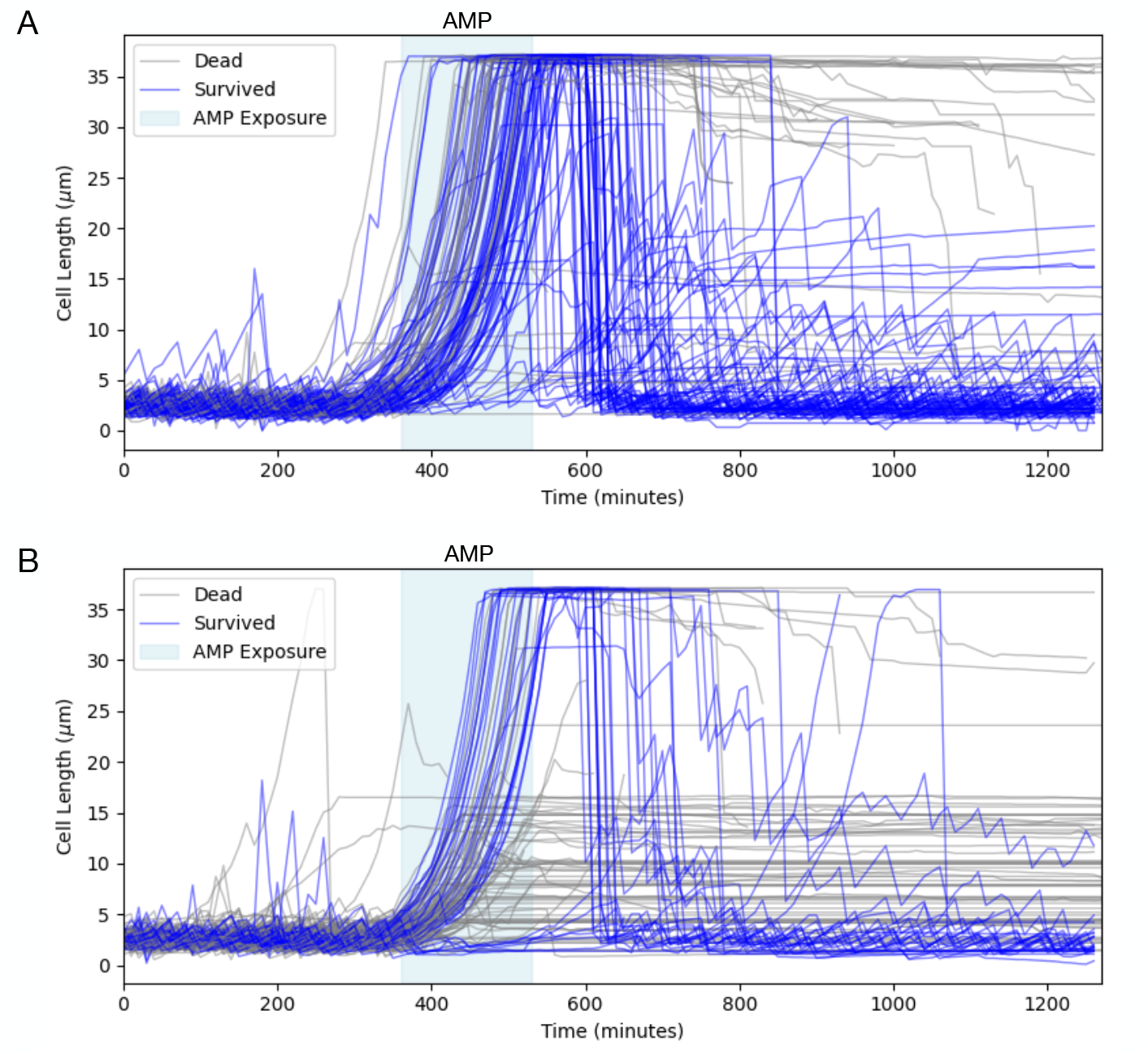
**A)** Time-series of individual cell lengths (*µ*m) for VIP205 cells tracked in the mother machine. The blue shaded area denotes the period of ampicillin (AMP) exposure. Cells that survived (resumed division after AMP removal) are shown in blue, while non-survivors are shown in gray. **B)** Time-series of individual cell lengths (*µ*m) for control (MC-SCFP3A) cells in the mother machine. The AMP exposure period is shaded in blue. Cells that survived are shown in blue, while those that did not are in gray.

**Figure S5.**
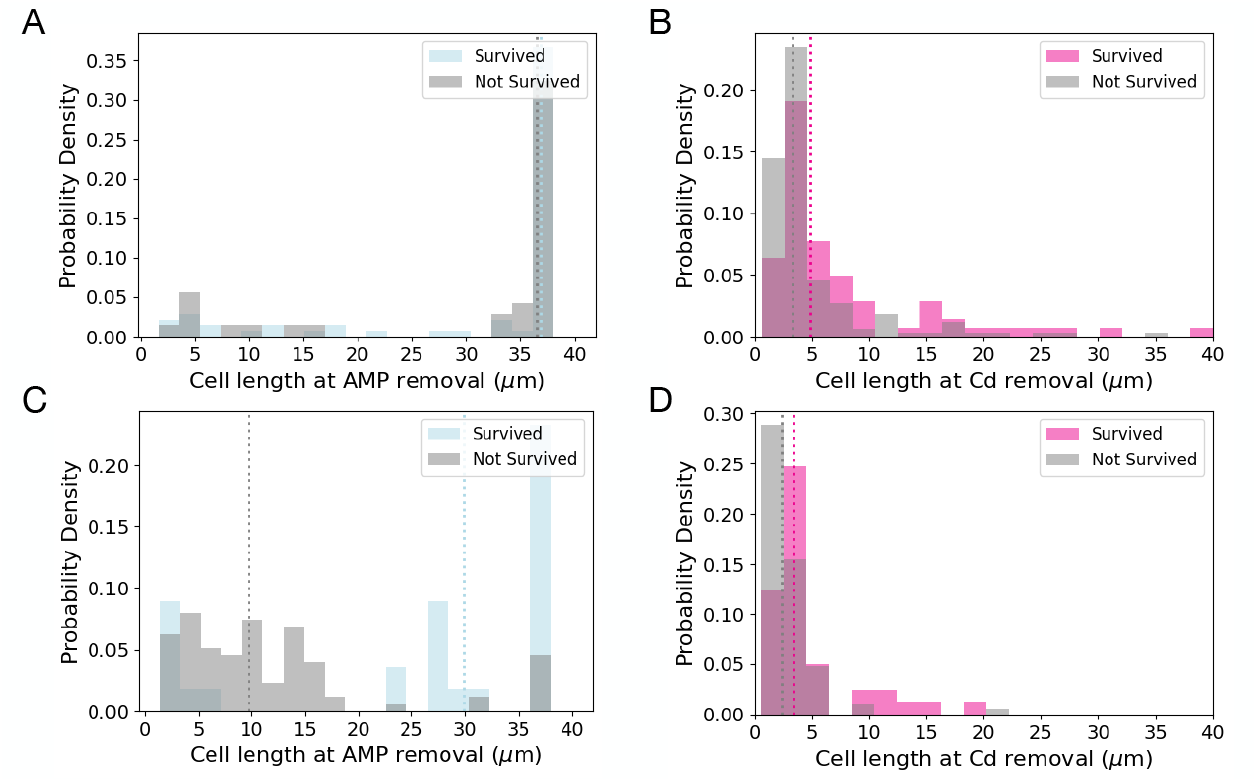
**A)** Probability density distribution of cell lengths at the moment of AMP exposure. Surviving cells are shown in blue, while non-survivors are shown in gray. **B)** Probability density distribution of cell lengths at the moment of Cd exposure. Surviving cells are shown in pink, while non-survivors are shown in gray. **C)** Cell length distribution for AMP-exposed populations, highlighting differences between survivors (blue) and non-survivors (gray). **D)** Cell length distribution for Cd-exposed populations, highlighting differences between survivors (pink) and non-survivors (gray).

**Figure S6.**
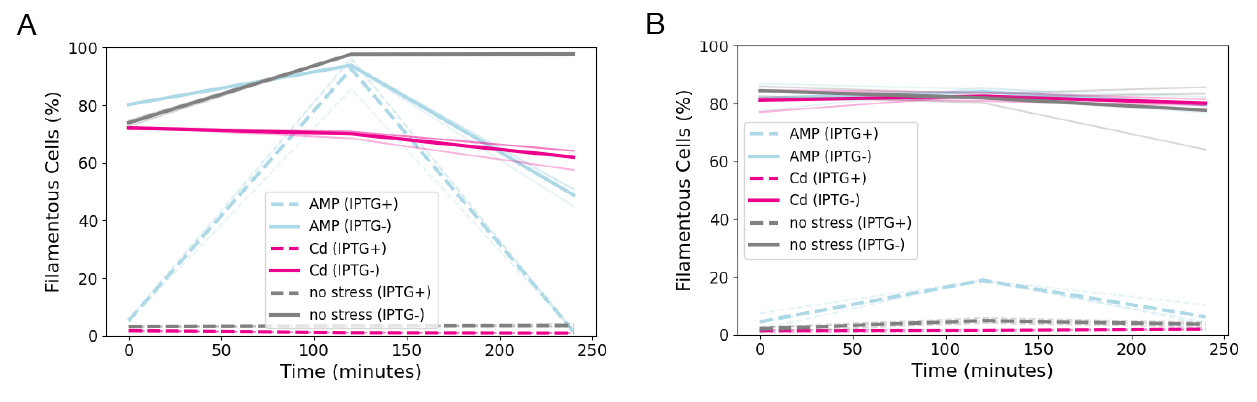
Filamentation dynamics in wild-type and ΔRecA strains under stress. **A)** Time-resolved quantification of filamentous cells in strain VIP205 exposed to AMP (blue), Cd (pink), or no stress (grey). Solid lines represent filamented populations (–IPTG) and dashed lines represent non-filamented controls (IPTG+). Thick lines show the mean across replicates; faint lines indicate individual experiments. B) Same as in A, but for VIP205 ΔRecA, which lacks the master regulator of the SOS response. Filamentation is still inducible with IPTG withdrawal, but AMP does not trigger endogenous filamentation in this strain.

**Figure S7.**
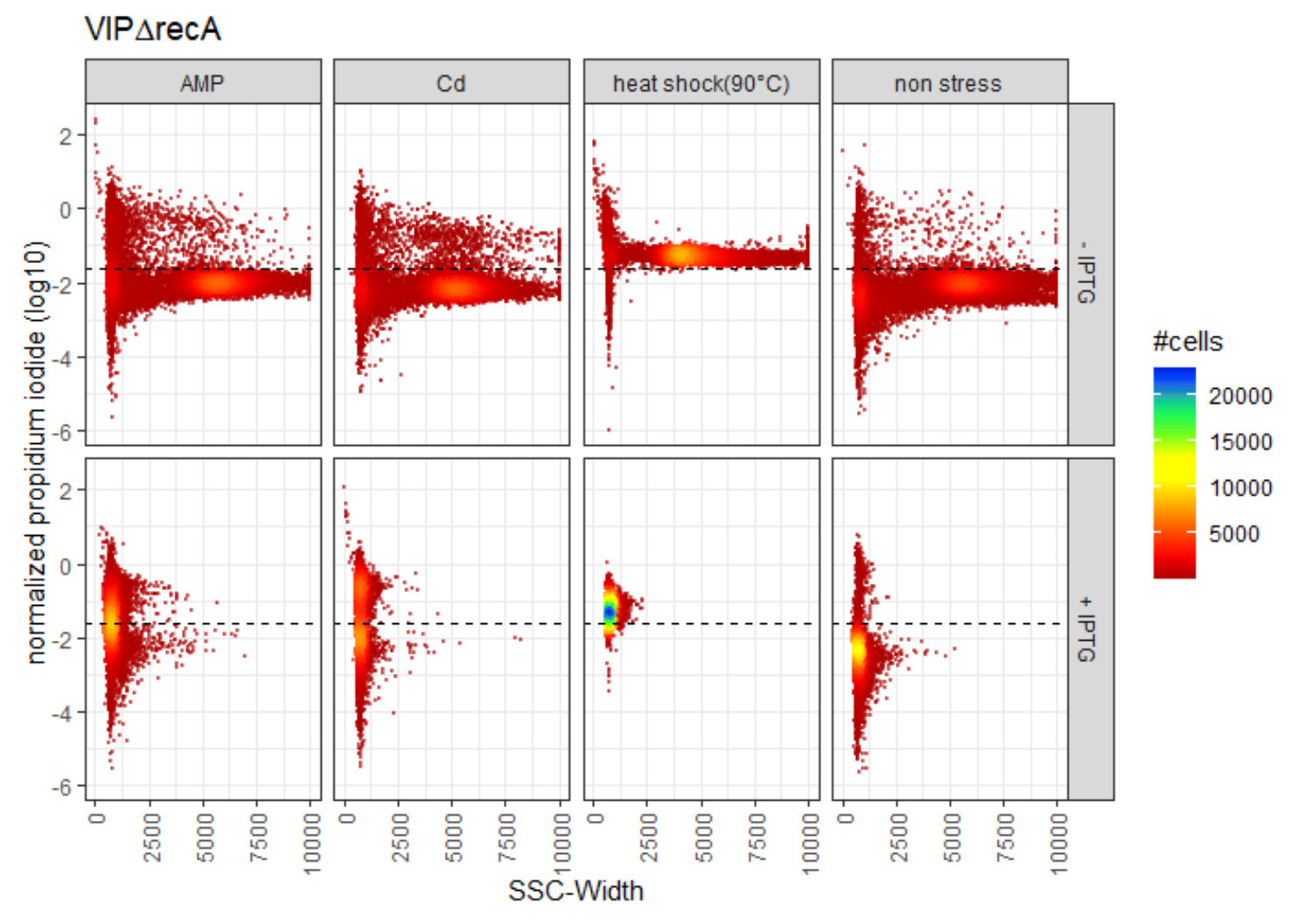
Filamentation and viability profiles of ΔRecA under multiple stress conditions. Flow cytometry distributions of SSC-Width (x-axis; proxy for cell filamentation) versus normalized propidium iodide signal (y-axis; loglO(FL2-A/SSC-A), proxy for cell death) in the ΔRecA strain. Top row: IPTG− (induced filamentation); Bottom row: IPTG+ (control, no induced filamentation). Columns represent different conditions: AMP, Cd, Heat Shock (90°C, positive control — all cells dead), and No Stress (negative control — all cells survive). Despite the inability to activate the SOS response, the induced system (IPTG−) can still produce filamented cells and confer protection under toxic stress. In contrast, IPTG+ results in higher death rates, both under AMP and Cd exposure.

**Table S1.** *Escherichia coli* strains used in this study.

| Strain | Genotype | Source |
| --- | --- | --- |
| MC1061 | F- $\Delta$(ara-leu)7697 [araD139]B/r $\Delta$(codB-lacI)3 galK16 galE15 λ- e14- mcrA0 relA1 rpsL150(StrR) spoT1 mcrB1 hsdR2(r-m+) | Gift from M. Vicente |
| VIP205 | MC1061, Ptac::ftsZ, KanR | Gift from M. Vicente |
| VIP205ΔRecA | VIP205, ΔRecA::Tn10 (TetR) | Gift from M. Vicente |
| MG1655 | F- λ-, rph-1 | Laboratory stock |
| MC-SCFP3A | MC1061, pEB1-SCFP3A:: KanR | This study |

**Table S2.** Parameters for numerical simulations.

| Parameter | Description | Value |
| --- | --- | --- |
| $L_0$ | Initial cell length | 10.0 $\mu\text{m}$ |
| $k_{in}$ | Antibiotic influx rate | 0.05 $\text{min}^{-1}$ |
| $k_L$ | Cell growth rate constant | 0.05 $\text{min}^{-1}$ |
| $r_{out}$ | Outer cell radius | 0.5 $\mu\text{m}$ |
| $r_{in}$ | Inner cell radius (periplasm) | 0.45 $\mu\text{m}$ |
| $A_{int,0}$ | Initial intracellular antibiotic concentration | 0.0 $\mu\text{g/mL}$ |
| $T_{\text{antibiotic}}$ | Time of antibiotic introduction | 20 min |
| $T_{\text{max}}$ | Total simulation time | 100 min |
| $k_{in,\text{mean}}$ | Mean antibiotic influx rate | 0.05 $\text{min}^{-1}$ |
| $k_{in,\text{std}}$ | Std. dev. antibiotic influx rate | 0.01 $\text{min}^{-1}$ |
| $A_{\text{ext}}$ | External antibiotic concentration range | 0–200 $\mu\text{g/mL}$ |
| $A_{int,0}$ | Initial intracellular antibiotic concentration | 0.0 $\mu\text{g/mL}$ |
| $A_{\text{lethal}}$ | Lethal intracellular antibiotic threshold | 20 $\mu\text{g/mL}$ |

